# Embedding covariate adjustments in tree-based automated machine learning for biomedical big data analyses

**DOI:** 10.1101/2020.08.24.265116

**Authors:** Elisabetta Manduchi, Weixuan Fu, Joseph D. Romano, Stefano Ruberto, Jason H. Moore

**Affiliations:** Department of Biostatistics, Epidemiology & Informatics, University of Pennsylvania, Philadelphia, PA 19104, USA; Institute for Biomedical Informatics, University of Pennsylvania, Philadelphia, PA 19104, USA

**Keywords:** AutoML, covariate adjustment, genetic programming, pathways, feature importance

## Abstract

**Background:** A typical task in bioinformatics consists of identifying which features are associated with a target outcome of interest and building a predictive model. Automated machine learning (AutoML) systems such as the Tree-based Pipeline Optimization Tool (TPOT) constitute an appealing approach to this end. However, in biomedical data, there are often baseline characteristics of the subjects in a study or batch effects that need to be adjusted for in order to better isolate the effects of the features of interest on the target. Thus, the ability to perform covariate adjustments becomes particularly important for applications of AutoML to biomedical big data analysis.

**Results:** We present an approach to adjust for covariates affecting features and/or target in TPOT. Our approach is based on regressing out the covariates in a manner that avoids ‘leakage’ during the cross-validation training procedure. We then describe applications of this approach to toxicogenomics and schizophrenia gene expression data sets. The TPOT extensions discussed in this work are available at https://github.com/EpistasisLab/tpot/tree/v0.11.1-resAdj.

**Conclusions:** In this work, we address an important need in the context of AutoML, which is particularly crucial for applications to bioinformatics and medical informatics, namely covariate adjustments. To this end we present a substantial extension of TPOT, a genetic programming based AutoML approach. We show the utility of this extension by applications to large toxicogenomics and differential gene expression data. The method is generally applicable in many other scenarios from the biomedical field.

## Background

Automated machine learning (AutoML) refers to methods which assist (potentially non-expert) users in the optimization of model construction steps such as data preprocessing, feature selection, feature transformations, model selection, and hyperparameter tuning.

The Tree-based Pipeline Optimization Tool (TPOT) [1, 2] is a genetic programming (GP) based AutoML which has been successfully used in biomedical applications including genetics [1], metabolomics [3, 4], and transcriptomics [5]. TPOT explores learning pipelines consisting of arbitrary combinations of selectors, transformers, and estimators (classifiers or regressors). In order to extend its scalability and provide more interpretable results, it is possible to specify a Template for the searched pipelines and to incorporate a Feature Set Selector (FSS) at the beginning of each pipeline to slice the input data set into smaller sets of features allowing the GP to select the best subset in the final pipeline [5].

A desirable feature of TPOT is the ability to adjust for relevant covariates, as this is particularly important in the biomedical context where there are often either baseline characteristics of the subjects or batch effects whose influence on the target or the features needs to be removed so to isolate the actual effects of the features on the target. It is important to note that, while common in biostatistics and epidemiology, covariate adjustment is uncommon and understudied in machine learning.

A typical covariate adjustment approach consists in ‘regressing the covariates out’ of the relevant variables in a data set, which could be the target, or some or all features or both (in case of confounding covariates). Essentially, to regress a set of covariates out of a variable, a predictor model for this variable based on such covariates is first built (e.g. using linear regression if the variable is continuous or logistic regression if it is binary or multiclass) and then the variable is replaced by the residuals from this model [6]. A simplified example of this can be found in [3]. However, the GP optimization in TPOT is based on cross-validation (CV). Consequently, simply regressing out the covariates from the relevant variables *before* feeding the data to TPOT suffers from ‘leakage’ because, for each CV split, a model built on the training part gets access to information it otherwise would not know, as it uses residuals calculated from a regression which employed the entire data set, i.e. both the training and the testing parts of the CV split. Such leakage can result in overfitting thus reducing the generalizability of the model.

In this work, we present an extension of the TPOT framework, referred to as ‘resAdj TPOT’ in what follows, which allows covariate adjustment without leakage. To illustrate the usefulness of this, we analyzed a toxicogenomic data set extracted from TG-GATEs [7] representing gene expression array data on kidney tissue from rats exposed to individual drugs known to cause kidney injury. Combining resAdj TPOT with the FSS and Template features, we identified pathways and genes whose expression is associated with creatinine levels in rat kidney, after removing confounding effects such as study batch, compound treatment, dose, and sacrifice time. Creatinine levels provide a broad image of overall kidney health and our findings are very consistent with known kidney biology. Moreover, if we apply TPOT with no covariate adjustments to this data set, the results are different and only tangentially associated with kidney disease. We also show an application to a gene expression dataset from PsychENCODE [8], where the selected covariates are not expected to bear much relevance. In this case we obtain similar results with resAdj and classic (i.e. no adjustments) TPOT, as desirable. The results of our comparisons between resAdj and classic TPOT confirm that our adjustment strategy has been useful in the first case and correctly neutral in the second. Interestingly, we also noted that in the second data set TPOT identified known pathways associated with differential expression between schizophrenic and controls which were not detected by the more typical Gene Set Enrichment Analysis (GSEA) [9].

## Methods

### TPOT leakage-free covariate adjustment

TPOT leverages the scikit-learn framework [10] and uses GP to evolve machine learning pipelines that consist of selectors, transformers and estimators [1, 2]. The GP seeks to optimize machine learning pipelines with respect to a specified score (e.g. ‘balanced accuracy’, ‘r-squared’, etc.) using CV to avoid overfitting on the provided data. Thus, in the pipeline optimization phase, the training set consists of a subset of the input samples (we used 75% at each CV split). At the end of the pipeline optimization procedure, the best pipeline is then trained on the entire set of input samples.

Suppose that, for each of *m* subjects, we have values for a target variable (binary, multiclass, or continuous) and a collection of *n* features (binary, multiclass, or continuous). We represent the target values by an *m-*dimensional vector ***y*** and the feature values by an *m×n* matrix ***X***. Suppose also that we have values for a collection of covariates that need to be adjusted for. Given a covariate, depending on the context of the data set, it may make sense to adjust only the target ***y*** by it, or only a subset of the features (columns in ***X)***, or both the target and a subset of the features (the latter is appropriate when the covariate is a confounder). We adjust the values of a variable *v* (which may be either the target or a feature) by a collection of covariates by ‘regressing the covariates out’, with the typical approach of fitting an estimator to *v* on the covariates (e.g. linear regression if *v* is continuous, or logistic regression otherwise) then replacing the values of *v* by the corresponding residuals. The latter are obtained by subtracting from the values of *v* the values predicted by the estimator (if *v* is continuous) or the expected values based on the estimator predicted class probabilities (if *v* is binary or multiclass). However, for each CV train-test split, to avoid leakage, *the estimator must be fitted only using the training data for that split* (see Additional file 1 for details). This can be easily achieved for feature covariate adjustments within the current TPOT/scikit-learn framework, but for target covariate (***y*)** adjustment it is necessary to substantially extend the framework.

For feature covariate adjustments, we have added a transformer (*resAdjTransformer*) to TPOT. This needs to be either the first step of any pipeline or the second step after an FSS (TPOT Template can be used to specify these). The initial input to TPOT adds the covariate columns to ***X***. One hyperparameter of this transformer is a file specifying which columns of ***X*** should be adjusted by which covariate columns. The transformer applies the no leakage residual adjustments to these columns and removes the covariate columns before passing its output on to the other steps. If no covariate adjustment on the target is needed, classic TPOT can then be run as usual (Figure 1 path A).

**Figure 1.**
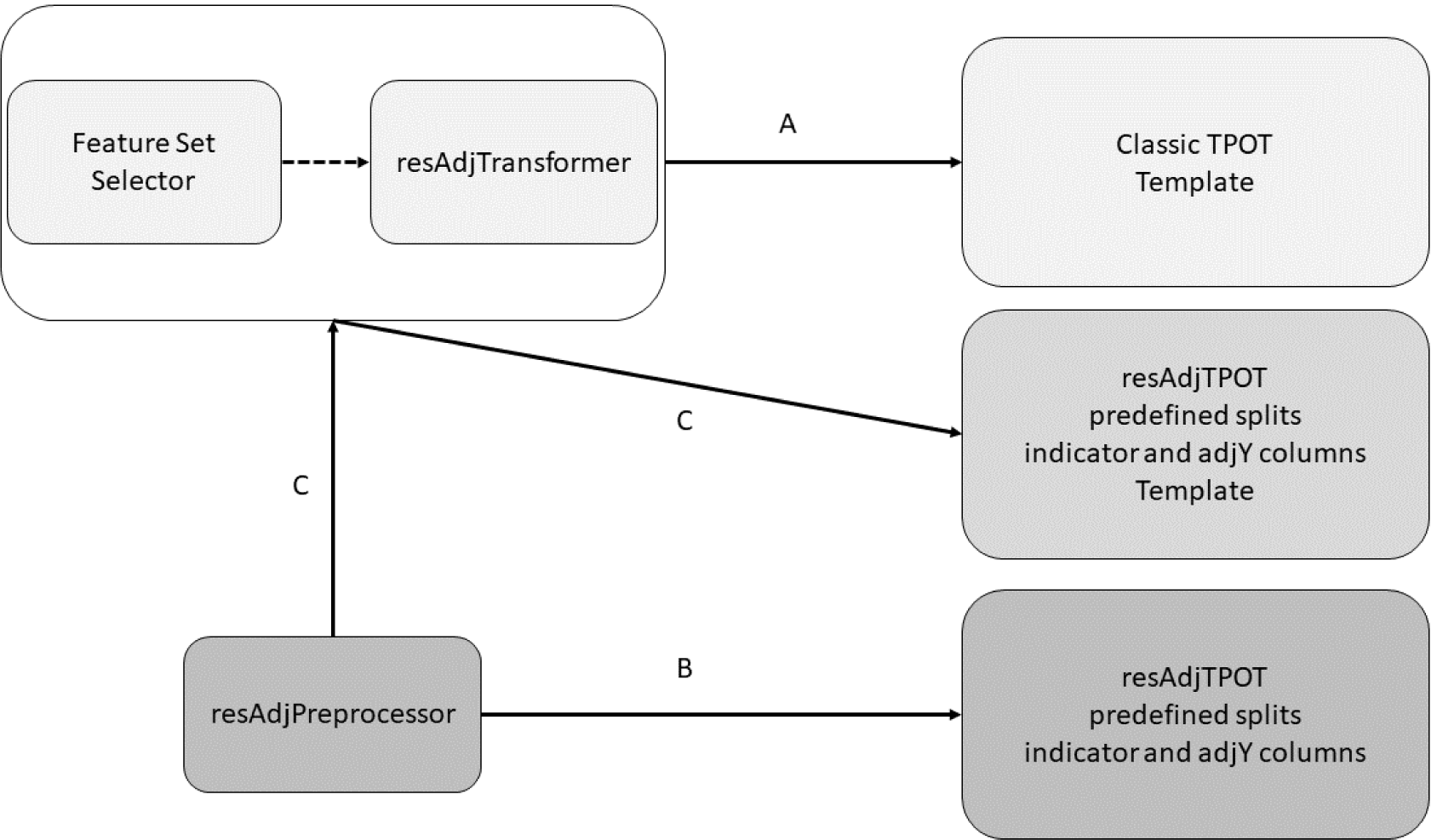
The 3 possible workflows for resAdj TPOT. (A) Workflow when the only needed adjustments are for features. The Feature Set Selector (FSS) step is optional. (B) Workflow when the only needed adjustments are for the target. adjY denotes the no-leakage adjustment of the target for each predefined CV split. (C) Workflow when adjustments are needed for both the target and features.

For target covariate adjustment, we have added a pre-processor, which creates predefined CV splits and, for each split, adds 2 columns to the dataset; an indicator column to denote the training-testing rows and a column with precomputed no-leakage residuals for ***y*** using that split. In addition, ***y*** is fully replaced by the residuals (this is only used for the final Testing Score reported by TPOT, after the optimal pipeline has been determined via CV). The output of this pre-processor (whose structure is outlined in Additional file 2) can then be passed on to TPOT, which must be run with the same CV splits. For each relevant scorer, selector, transformer, and estimator in the classic TPOT, we have added a corresponding scorer, selector, transformer, and estimator which, by using the indicator and corresponding residual columns, enables the pipeline to flow utilizing the appropriately adjusted target at each CV split. The pre-processor can optionally also prepare a hold-out testing set to assess the optimized pipelines output by resAdj TPOT runs.

Note that, when the only needed adjustments are for features, classic TPOT can be run as long as the *resAdjTransformer* is incorporated within every pipeline (using Template, see Figure 1 path A). If the only needed adjustment is for the target, then the pre-processor and the new scorer, selectors, transformers, and estimators must be used with predefined CV splits (Figure 1 path B). If both features and target need adjustment, then all the above can be used together (Figure 1 path C).

## Data sets

### TG-GATEs

We selected from TG-GATEs [7] the 933 microarray gene expression assays on kidney tissue from rats in the interquartile range of kidney weight, where each rat was treated with one of 41 compounds in a single dose. The dose was one of ‘control’ (untreated animal), ‘low’, ‘medium’, or ‘high’ (concentrations varied based on the compound) and the rat was sacrificed after a duration of 3, 6, 9 or 24 hours. We chose creatinine level measured at sacrifice time as the target of interest, because this provides a broad image of overall kidney health. After obtaining the CEL files from accession E-MTAB-799 in Array Express [11], we summarized and normalized (with RMA) the expression values using the Transcriptome Analysis Console (TAC; Affymetrix) software. We encoded dose and sacrifice time by ordinal variables with 4 levels and creatinine measurements by an ordinal variable with 7 levels. We encoded the 41 compounds with 6 binary variables using the binary encoder from the Category-Encoders (http://contrib.scikit-learn.org/category_encoders/). We defined 154 Feature Sets corresponding to the pathways obtained by mapping to rat the MSigDB [12] canonical pathways from KEGG and the pathways annotated in the Rat 230 2.0 Affymetrix Array. The features in our analyses were the 1632 probesets mapping to these Feature Sets.

### PsychENCODE

We downloaded the prefrontal cortex normalized RNAseq gene expression data from http://resource.psychencode.org/Datasets/Derived/DER-01_PEC_Gene_expression_matrix_normalized.txt. We extracted 1072 control and schizophrenic individuals with either Caucasian or African American ethnicity, from 3 studies (LIBD_szControl, CMC, and BrainGVEX). Disease status was our target. We encoded each of disease status, sex and ethnicity by binary variables and study by 2 binary variables (using one-hot encoding). We defined 186 Feature Sets corresponding to the MSigDB canonical pathways from KEGG. The features in our analyses were the 4952 genes mapping to these Feature Sets.

## Results

### TG-GATEs

To fully exploit this large expression data set to identify pathways and genes directly associated to creatinine levels, we needed to factor out the confounding effect of compound treatment. To explore the latter, we clustered the expression data from the 933 assays using *k*-means [13, 14]. We leveraged NbClust [15] to identify the value of *k* (between 30 and 200) maximizing the Dunn index [16], a measure of cluster quality defined as the ratio between minimal intercluster distance to maximal intracluster distance. The best Dunn index was 0.52 for *k*=171. We then measured the Biological Homogeneity Index (BHI) of the resulting clustering with respect to various annotations, where the BHI (defined in https://cran.r-project.org/web/packages/clValid/vignettes/clValid.pdf) is a value in the range [0, 1], reflecting the average proportion of pairs in the same cluster with identical annotation. Thus, larger BHI values correspond to more homogeneous clusters in terms of the annotation attribute. We calculated the BHI for each of compound, dose, and sacrifice time. The values were 0.77, 0.31, and 0.66, which indicates that these are important covariates to adjust for, as they are associated to how the data are clustering. We also note that, since TG-GATEs combines data collected from different studies through the years, adjusting all features and target by compound has the additional benefit of removing study batch effects, as typically studies revolved around specific compounds.

To assess the robustness of TPOT with respect to both GP stochasticity and selection of training and testing portions, it is crucial to run the program repeatedly with different random train/test splits and verify the consistency of the results across such runs, in particular in terms of selected pathways. We generated 100 random splits of the data into training (75%) and testing (25%) parts. For each split, we ran resAdj TPOT, adjusting target and all features by the encoded compounds, doses, and sacrifice times (following path C in Figure 1), using 500 generations and a population of 500 in the GP. For each run, the training data set underwent 5 CV splits and the adjustments during training were leakage-free thanks to the approach we described in Methods. Since the target is replaced by residuals, it represents a continuous outcome. We used the Template ‘FeatureSetSelector-resAdjTransformer-Transformer-Regressor’ and the coefficient of determination (R^2^) to score the models. For each run we noted the optimal pipeline, including the selected pathway (i.e. Feature Set) and its score on the held-out testing set. Figure 2 summarizes the results. The ‘G-protein signaling’ and ‘Integrin-mediated cell adhesion’ pathways were consistently selected in most of the runs (54% and 28%, respectively) with a median R^2^ of ∼0.57. Both pathways play known roles in kidney function: G-protein signaling is responsible for mediating response to various types of physical damage to the cell in a broad range of renal diseases [17, 18]. Integrin-mediated cell adhesion plays a crucial structural role in the portions of the kidneys responsible for collecting waste from the circulatory system [19]. Therefore, considering that creatinine is an indicator of kidney health and is not restricted to a specific mechanism of kidney injury, when adjusting for relevant covariates, TPOT preferentially selects biologically plausible pathways.

**Figure 2.**
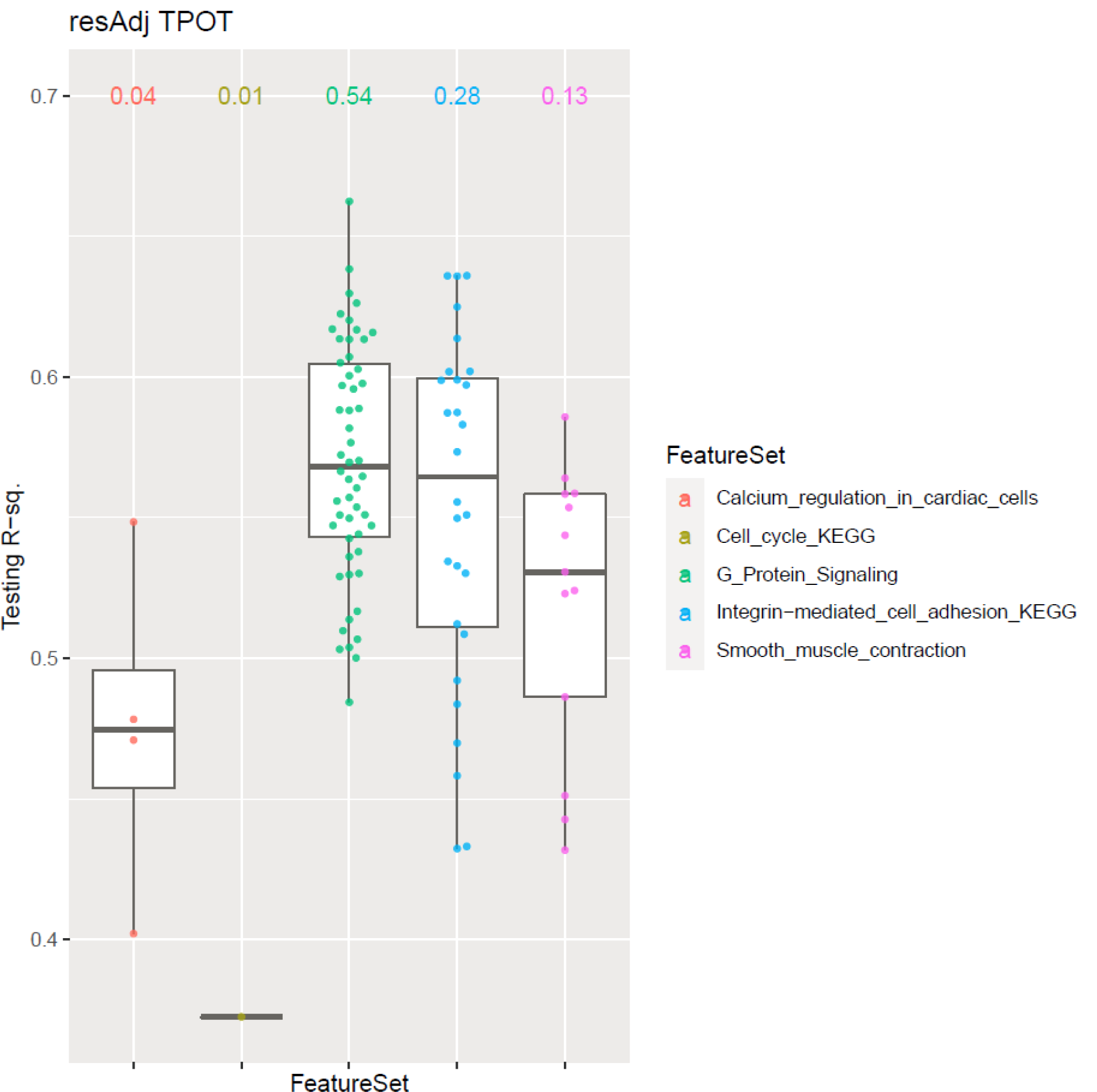
Boxplots for the results of 100 runs of resAdj TPOT on the TG-GATEs data set. Each point corresponds to one run, where the *x*-coordinate indicates the pathway (Feature Set) selected in the optimal pipeline for that run and the *y*-coordinate indicates the R^2^ on the held-out Testing dataset.

We also assessed features and covariates by calculating their permutation importance using eli5 v0.10.1 (https://github.com/TeamHG-Memex/eli5) across the 100 runs and then computing the weighted (by testing score) average of the mean score decrease as a percentage of the score. As illustrated in Figure 3, the adjustments by compound and sacrifice time are highly relevant. Moreover, the top 20 features (ranked by importance scores) include genes that are biologically significant in terms of modulating kidney function. For example, the gene with the highest importance score, *Akap9*, codes A-Kinase Anchoring Protein 9, which is highly expressed in kidney tissue, is implicated in retention of T-lymphocytes in kidney tissue under states of inflammation [20], and is one of the most commonly mutated genes in metastatic renal carcinoma [21]. The gene *Gnb1*, which codes for G Protein Subunit β 1, comprises both the second and third most important features (at two genetic loci), and is highly expressed in kidney glomeruli and tubules [22]. Like *Akap9, Gnb1* is also implicated in kidney disease, where downregulation of the gene is associated with worsened prognosis of clear-cell renal cell carcinoma [23], as well as in resistance to tyrosine kinase inhibitor drugs in human models of kidney cancer [24]. Similar patterns can be seen in other important features, although we omit them for brevity.

**Figure 3.**
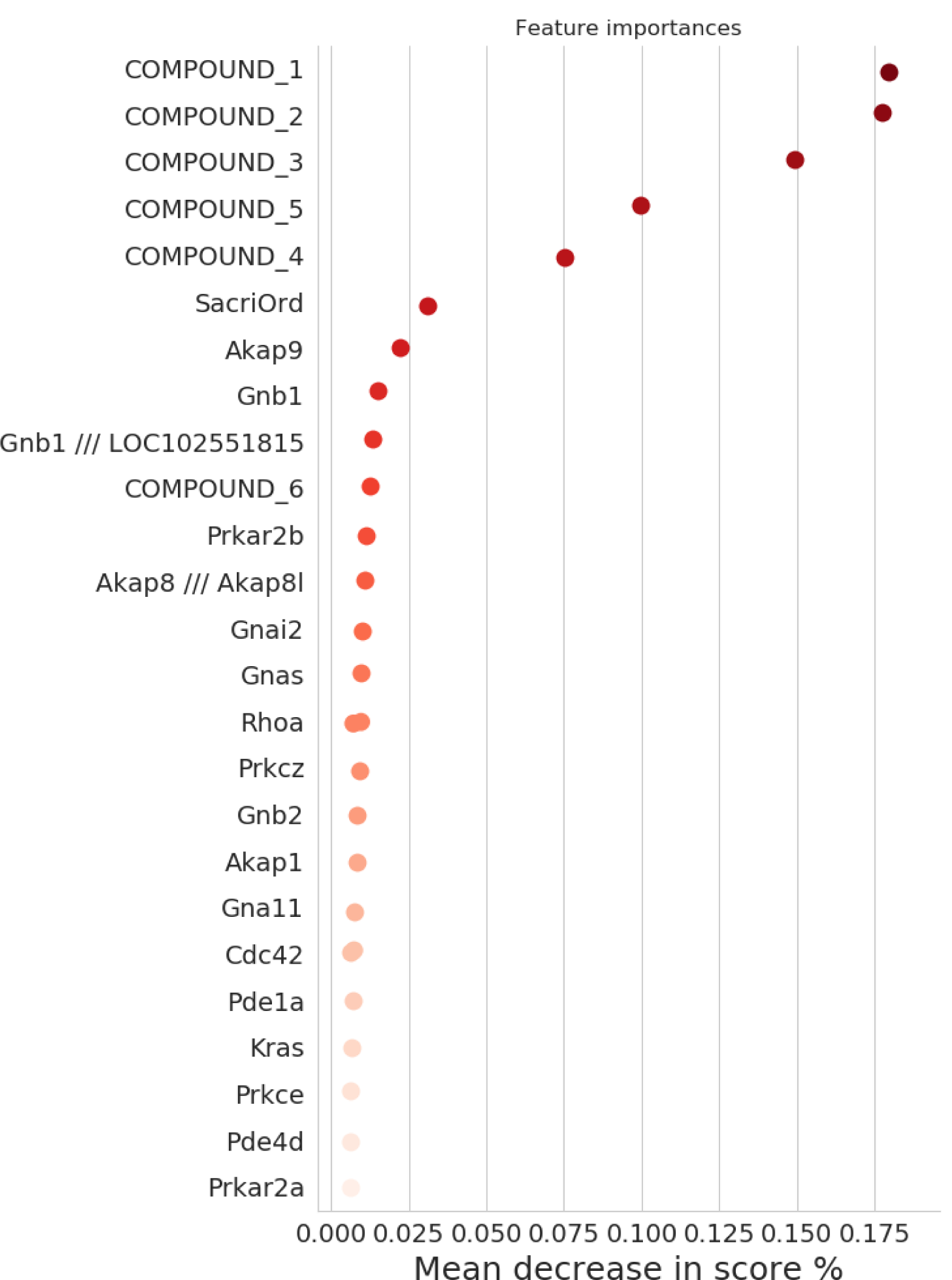
Permutation importance from 100 runs of resAdj TPOT on the TG-GATEs data set. The top 20 features and 7 covariates are shown. The gene names are displayed on the *y*-axis and the weighted (by testing score) averages of the mean score decrease as a percentage of the score are displayed on the *x*-axis. The analyses were done at the probeset level, and for *Rhoa* there were two probesets among the top 20 features.

The clustering results underscore how adjusting for covariates in this data set is crucial to properly examine the net effect of genes on creatinine levels. Indeed, running classic TPOT (i.e. no covariate adjustments) on the expression features, yields quite different results. We used the same 100 random splits into training and testing parts and, for each split, we ran TPOT with the template ‘FeatureSetSelector-Transformer-Classifier’ and balanced accuracy as a scorer since, without adjustments, the target is the original multiclass variable (7 levels of creatinine). Again, we used 500 generations and a population of 500 in the GP. As illustrated in Additional file 3, the most frequently selected pathway was ‘Smooth-muscle contraction’ (45% of the runs) followed by ‘mRNA-processing’ (14%). Parenthetically, scores cannot be directly compared with the runs in resAdj TPOT since the scorers are different (balanced accuracy versus R^2^). Even though both pathways that were identified as important in the resAdj runs (‘G-protein signaling’ and ‘Integrin-mediated cell adhesion’) also show up in some of the classic runs, they are partially obscured by a number of other pathways that are largely uninformative or irrelevant in terms of kidney disease, primarily pathways that mediate the functionality of the heart and blood vessels. Although these are peripherally involved in kidney function due to the kidneys’ role in the circulatory system, there are no obvious connections between these pathways and kidney injury in particular.

In terms of feature importance, the difference between classic and resAdj TPOT is striking. Each of the top two most important genes in the covariate adjusted runs (*Akap9* and *Gnb1*, but not the second locus of *Gnb1*) are notably absent from the list of the top 20 features from the unadjusted runs (see Additional file 4), implying that the covariate adjustment filters out spurious correlations that would otherwise obscure mechanistically significant features, at least when applied to data sets with a focus on toxicogenomic response. Rather, the top gene from the classic runs is *Sf3b1*, which is only tangentially associated with kidney disease via copy number variation, through no known mechanism [25].

### 3.2. PsychENCODE

In this data set, we considered sex and ethnicity as potential covariates affecting the target (disease status), and ethnicity and study as potential covariates affecting the features (measured gene expression). We therefore ran similar clustering analyses and BHI calculations to those done on TG-GATEs. Varying *k* from 10 to 240 for clustering of the 1072 expression assays, the best Dunn index was 0.47 corresponding to *k*=201. The BHI values of the corresponding clustering for the potential covariates affecting gene expression were 0.04 for both ethnicity and study, close to 0 and much smaller than the covariates we analyzed in TG-GATEs, indicating that these covariates do not have a strong effect on the expression. Thus, on these data, it is desirable to get similar results when running resAdj and classic TPOT. Indeed, this was the case. We proceeded similarly to TG-GATEs to set up 100 runs for each of resAdj and classic TPOT, noting the pathways identified in the optimal pipelines and computing permutation importance scores for the genes (see Additional files 5-8 for corresponding plots). In either classic or resAdj modality, for about 80% of the runs the pathway from the optimal pipeline was ‘Calcium Signaling’. The latter is an important pathway for the pathophysiology of schizophrenia, as implicated by several previous studies [26–31]. This pathway contained the top strongest genes based on permutation importance in both TPOT modalities, namely *PTAFR, GNAL, CAMK2G*, and *PRKCG*. The remaining runs, in both modalities, identified three other pathways whose association with schizophrenia is supported in the literature: ‘neuroactive ligand-receptor interaction’ [27, 30, 32], ‘MAPK signaling’ [33–35], and ‘pathways in cancer’ [35]. In addition to these, one of the classic runs identified ‘Long-Term potentiation’ (albeit with a lower score), which too has been reported as associated with schizophrenia [36]. We note that none of these 5 pathways were identified when we ran the typical GSEA [9] analyses on this data set. GSEA did not identify any of the MSigDB canonical pathways as significant at FDR<25%. Additional file 9 lists the pathways with GSEA unadjusted p-values < 0.05.

## Discussion

An important need for applications of AutoML methods, and machine learning in general, to biomedical data analyses is the ability to incorporate covariate adjustments. In this work, we have presented an approach to adjust the target and/or any subset of the features by a collection of relevant covariates in the context of TPOT, a GP-based AutoML approach. Our method enables adjustments while properly avoiding leakage during pipeline training. We have illustrated its usefulness by an application to a toxicogenomic dataset where adjustments were necessary in order to identify pathways and genes associated to creatinine levels in the presence of confounders. In this application we obtained results which were robust (across 100 runs) and very consistent with known kidney biology. We then applied resAdj TPOT to a different gene expression data set (schizophrenia), obtaining results which were extremely stable and supported by biological literature. In the latter data set we had an indication that confounding was less of an issue, and indeed we observed that resAdj led to the same results as classic TPOT, as desired. In general, though, selecting which covariates are important for adjusting which of the features or target is a delicate step, as different choices could significantly impact the results. Thus, developing approaches and tools aimed at assisting users in this endeavor is as an important area of research that needs to move in parallel with the refinement of approaches like ours.

Another area that needs to be further explored is how to best encode nominal variables (i.e. categorical variables with no natural order relationship) in the context of our method. When the number of values *N* of a nominal variable is small, one-hot encoding can be a simple solution. In this case, the variable is replaced by *N-1* new binary ‘dummy variables’. However, for large values of *N*, this encoding may not work well. In fact, when we tried it for the compound variable in our TG-GATEs data set, we obtained models with very low R^2^ (data not shown). In our case, switching to a binary encoding with six variables led to good results with high testing scores. But a systematic study of the effects of different encoding choices on our method is an important aspect for future research.

Our no-leakage adjustments leverage the typical approach of replacing a variable by its residuals obtained by regressing out the covariates via linear or logistic regression, depending on the type of variable. However, in principle, other estimators could be used, i.e. other classifiers for binary or multiclass variables and other regressors for continuous variables. Possibly, a first pass assessment of the best estimators to use for the adjustments could be done using TPOT itself. Refining and extending resAdj TPOT to this end is another interesting path for future developments.

## Conclusions

Our resAdj TPOT approach represents a first step towards addressing a relevant need for AutoML applicability to biomedical big data analyses, where covariate adjustments are often necessary. The applications we presented in this work leveraged toxicogenomics and differential gene expression data. But there are many other scenarios where resAdj TPOT could provide a useful analysis option. For example, it could aid in epistasis analyses of genotype data, where covariate adjustments (e.g. genetic principal components, age, sex, etc.) is typically crucial.

More generally, the increased availability of resources such as the UK Biobank [37] providing a rich and large basket of phenotypes and genotypes, enables a plethora of interesting data explorations for which methods such as resAdj TPOT represent a very useful tool. We anticipate this work will have broader impact on machine learning.

## List of abbreviations

AutoML: Automated Machine Learning
CV: Cross Validation
FSS: Feature Set Selector
GP: Genetic Programming
GSEA: Gene Set Enrichment Analysis
resAdj: residual adjustment
TPOT: Tree-based Pipeline Optimization Tool

## Declarations

### Availability of data and materials

We originally obtained the TG-GATEs CEL files used in this study from https://www.ebi.ac.uk/arrayexpress/experiments/E-MTAB-799. These files are currently available from ftp://ftp.biosciencedbc.jp/archive/open-tggates/LATEST/Rat/in_vivo/Kidney/Single/. The complete sample annotation can be obtained from ftp://ftp.biosciencedbc.jp/archive/open-tggates/LATEST/Open-tggates_AllAttribute.zip.

The list of 933 assays we used for our analyses is in Additional file 10. The PsychENCODE data can be obtained from http://resource.psychencode.org/.

The TPOT extensions discussed in this work are available at https://github.com/EpistasisLab/tpot/tree/v0.11.1-resAdj.

### Authors’ Contributions

EM conceived the TPOT extension, provided code, searched for data sets, analyzed the data, and drafted the manuscript. WF worked on the code and github repository and contributed to method discussions. JDR suggested the TG-GATEs data set, summarized the CEL files, and provided biological interpretations of the TG-GATEs results. SR contributed to several method discussions. JHM supervised the project and contributed to method discussions. All authors reviewed and edited the manuscript. All authors read and approved the final manuscript.

## Acknowledgments

The PsychENCODE data used to illustrate an application of our method were generated as part of the PsychENCODE Consortium supported by: U01MH103339, U01MH103365, U01MH103392, U01MH103340, U01MH103346, R01MH105472, R01MH094714, R01MH105898, R21MH102791, R21MH105881, R21MH103877, and P50MH106934 awarded to: Schahram Akbarian (Icahn School of Medicine at Mount Sinai), Gregory Crawford (Duke), Stella Dracheva (Icahn School of Medicine at Mount Sinai), Peggy Farnham (USC), Mark Gerstein (Yale), Daniel Geschwind (UCLA), Thomas M. Hyde (LIBD), Andrew Jaffe (LIBD), James A. Knowles (USC), Chunyu Liu (UIC), Dalila Pinto (Icahn School of Medicine at Mount Sinai), Nenad Sestan (Yale), Pamela Sklar (Icahn School of Medicine at Mount Sinai), Matthew State (UCSF), Patrick Sullivan (UNC), Flora Vaccarino (Yale), Sherman Weissman (Yale), Kevin White (UChicago) and Peter Zandi (JHU).

## Additional files

**Additional file 1. *Additional methods***. Details for the leakage-free covariate adjustment. (AdditionalFile1.pdf).

**Additional file 2. *resAdj TPOT pre-processor output***. Structure and column description of the output from this pre-processor (AdditionalFile2.pptx).

**Additional file 3. *Boxplots for the results of 100 runs of classic TPOT on the TG-GATEs data set***. Each point corresponds to one run, where the *x*-coordinate indicates the pathway (Feature Set) selected in the optimal pipeline for that run and the *y*-coordinate indicates the balanced accuracy on the held-out Testing dataset. (AdditionalFile3.pdf).

**Additional file 4. *Permutation importance from 100 runs of classic TPOT on the TG-GATEs data set***. The top 20 features are shown. The gene names are displayed on the *y*-axis and the weighted (by testing score) averages of the mean score decrease as a percentage of the score are displayed on the *x*-axis. The analyses were done at the probeset level, and for some of the genes there were two probesets among the top 20 features. (AdditionalFile4.png).

**Additional file 5. *Boxplots for the results of 100 runs of resAdj TPOT on the PsychENCODE data set***. Each point corresponds to one run, where the *x*-coordinate indicates the pathway (Feature Set) selected in the optimal pipeline for that run and the *y*-coordinate indicates the R^2^ on the held-out Testing dataset. (AdditionalFile5.pdf).

**Additional file 6. *Permutation importance from 100 runs of resAdj TPOT on the PscychENCODE data set***. The top 20 genes (features) and 2 covariates are shown. The gene names are displayed on the *y*-axis and the weighted (by testing score) averages of the mean score decrease as a percentage of the score are displayed on the *x*-axis. (AdditionalFile6.png).

**Additional file 7. *Boxplots for the results of 100 runs of classic TPOT on the PsychENCODE data set***. Each point corresponds to one run, where the *x*-coordinate indicates the pathway (Feature Set) selected in the optimal pipeline for that run and the *y*-coordinate indicates the balanced accuracy on the held-out Testing dataset. (AdditionalFile7.pdf).

**Additional file 8. *Permutation importance from 100 runs of classic TPOT on the PscychENCODE data set***. The top 20 genes (features) are shown. The gene names are displayed on the *y*-axis and the weighted (by testing score) averages of the mean score decrease as a percentage of the score are displayed on the *x*-axis. (AdditionalFile8.png).

**Additional file 9. *GSEA results on the PsychENCODE data set***. Pathways with unadjusted p-values<0.05. (AdditionalFile9.xlsx).

**Additional file 10. *TG-GATEs assays used***. The list of 933 assays used in this work (AdditionalFile10.txt).

