## Additional file 1 for "Embedding covariate adjustments in tree-based automated machine learning for biomedical big data analyses"

### Leakage-free covariate adjustment

Let $v$ be the target variable or feature which we want to adjust by covariates $C_1$, $C_2$, ..., $C_r$. Denote by $v_i$ and $C_{i1}$, $C_{i2}$, ..., $C_{ir}$ the observed values of $v$ and the $C$s for subject $i$.

Suppose we are given a cross-validation split of the subjects into a training and a testing set.

#### Continuous $v$

If $v$ is continuous, we use the **training set** to fit a linear regression model:

$$v = \beta_0 + \beta_1 C_1 + \beta_2 C_2 + \ldots + \beta_r C_r.$$

For each subject $i$ **(whether in the training or testing set)**, we subtract the model-fitted values from $v_i$:

$$v_i^{\text{adj}} = v_i - (\hat{\beta}_0 + \hat{\beta}_1 C_{i1} + \hat{\beta}_2 C_{i2} + \ldots + \hat{\beta}_r C_{ir}).$$

#### Binary $v$

If $v$ is binary with values are 0 and 1, we use the **training set** to fit a logistic regression model:

$$\text{logit}(\pi) = \beta_0 + \beta_1 C_1 + \beta_2 C_2 + \ldots + \beta_r C_r$$

where $\pi = Prob(v = 1)$.

For each subject $i$ **(whether in the training or testing set)**, we subtract the model-fitted values from the observed outcomes:

$$v_i^{\text{adj}} = v_i - \hat{\pi}_i$$

where

$$\hat{\pi}_i = \frac{\exp(\hat{\beta}_0 + \hat{\beta}_1 C_{i1} + \hat{\beta}_2 C_{i2} + \ldots + \hat{\beta}_r C_{ir}}{1 + \exp(\hat{\beta}_0 + \hat{\beta}_1 C_{i1} + \hat{\beta}_2 C_{i2} + \ldots + \hat{\beta}_r C_{ir})}.$$

#### Multiclass $v$

This is a simple extension of the binary case. If $v$ is multiclass with values $0, 1, \ldots, K$, we use the **training set** to fit a multinomial logistic regression model, deriving values $\hat{\pi}_k$ for $k = 1, \ldots, K$.

For each subject $i$ **(whether in the training or testing set)**, we subtract the model-fitted values from the observed outcomes:

$$v_i^{\text{adj}} = v_i - \sum_{k=1}^{K} \hat{\pi}_{ki} k.$$
