## Additional file 2 for "Embedding covariate adjustments in tree-based automated machine learning for biomedical big data analyses"

### Slide 1
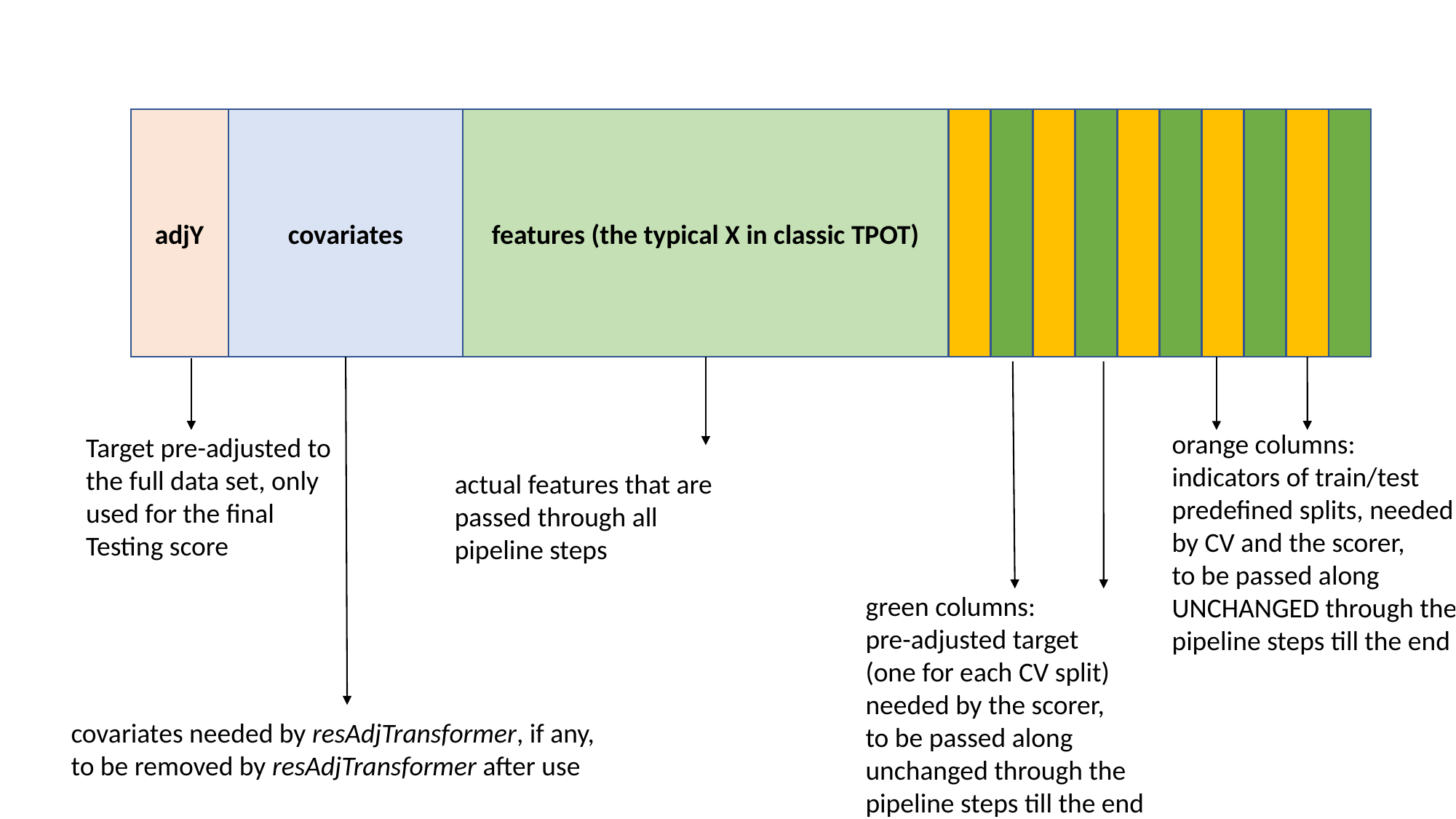

adjY
covariates
features (the typical X in classic TPOT)
orange columns:
indicators of train/test
predefined splits, needed
by CV and the scorer,
to be passed along
UNCHANGED through the
pipeline steps till the end
Target pre-adjusted to
the full data set, only
used for the final
Testing score
actual features that are
passed through all
pipeline steps
green columns:
pre-adjusted target
(one for each CV split)
needed by the scorer,
to be passed along
unchanged through the
pipeline steps till the end
covariates needed by resAdjTransformer, if any,
to be removed by resAdjTransformer after use
