## Supplementary figures and images for "Embedding covariate adjustments in tree-based automated machine learning for biomedical big data analyses"

### Additional file 3

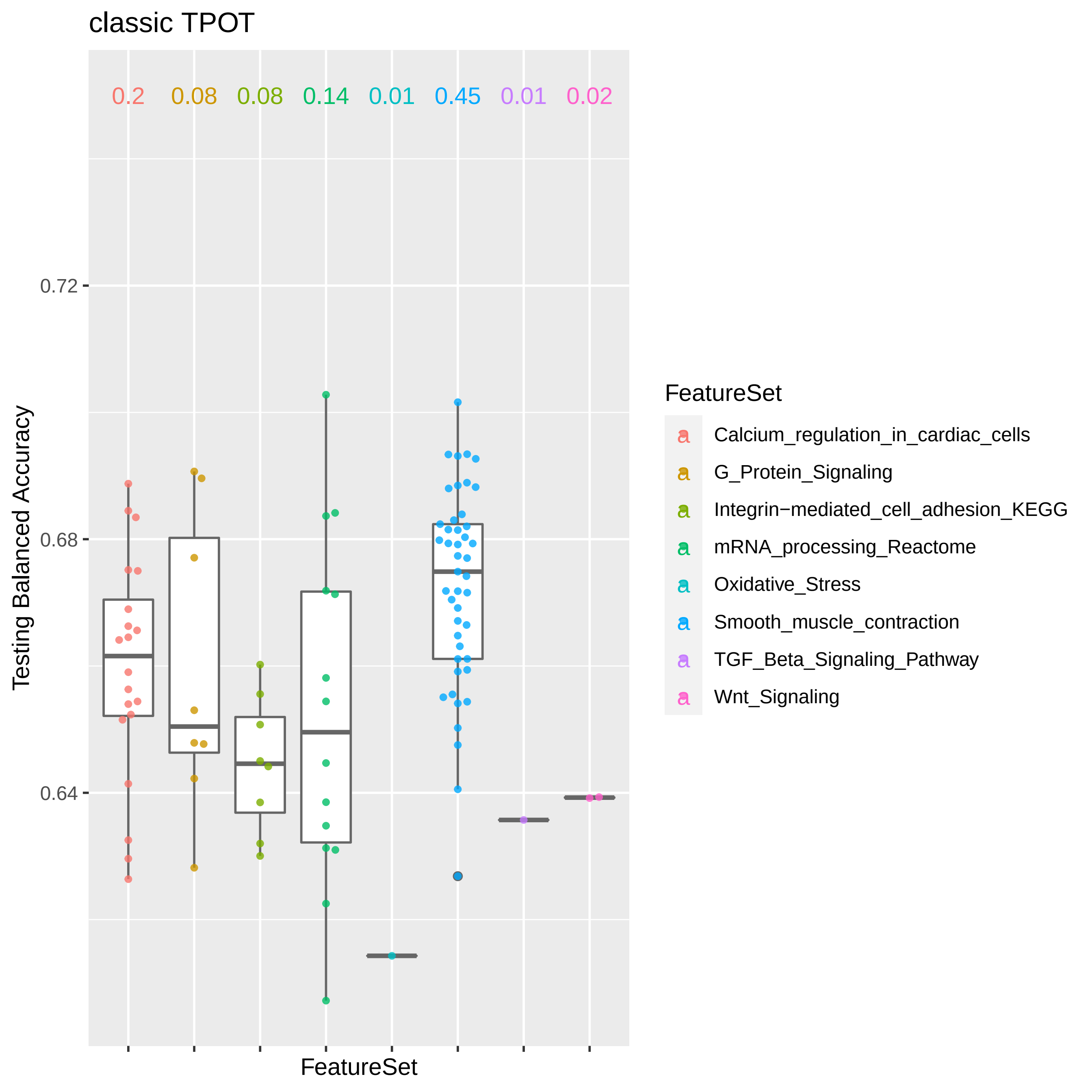

### Additional file 4

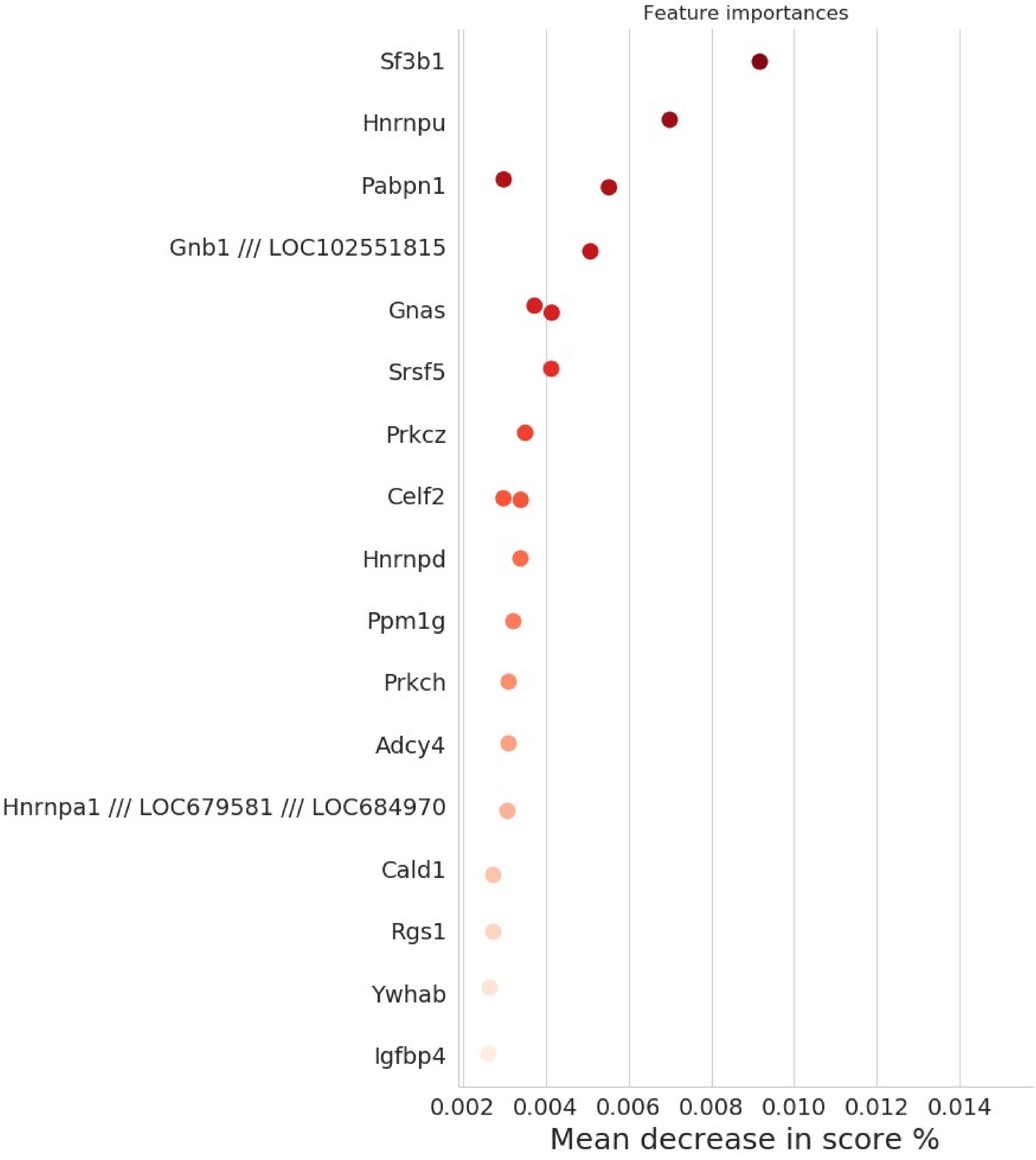

### Additional file 5

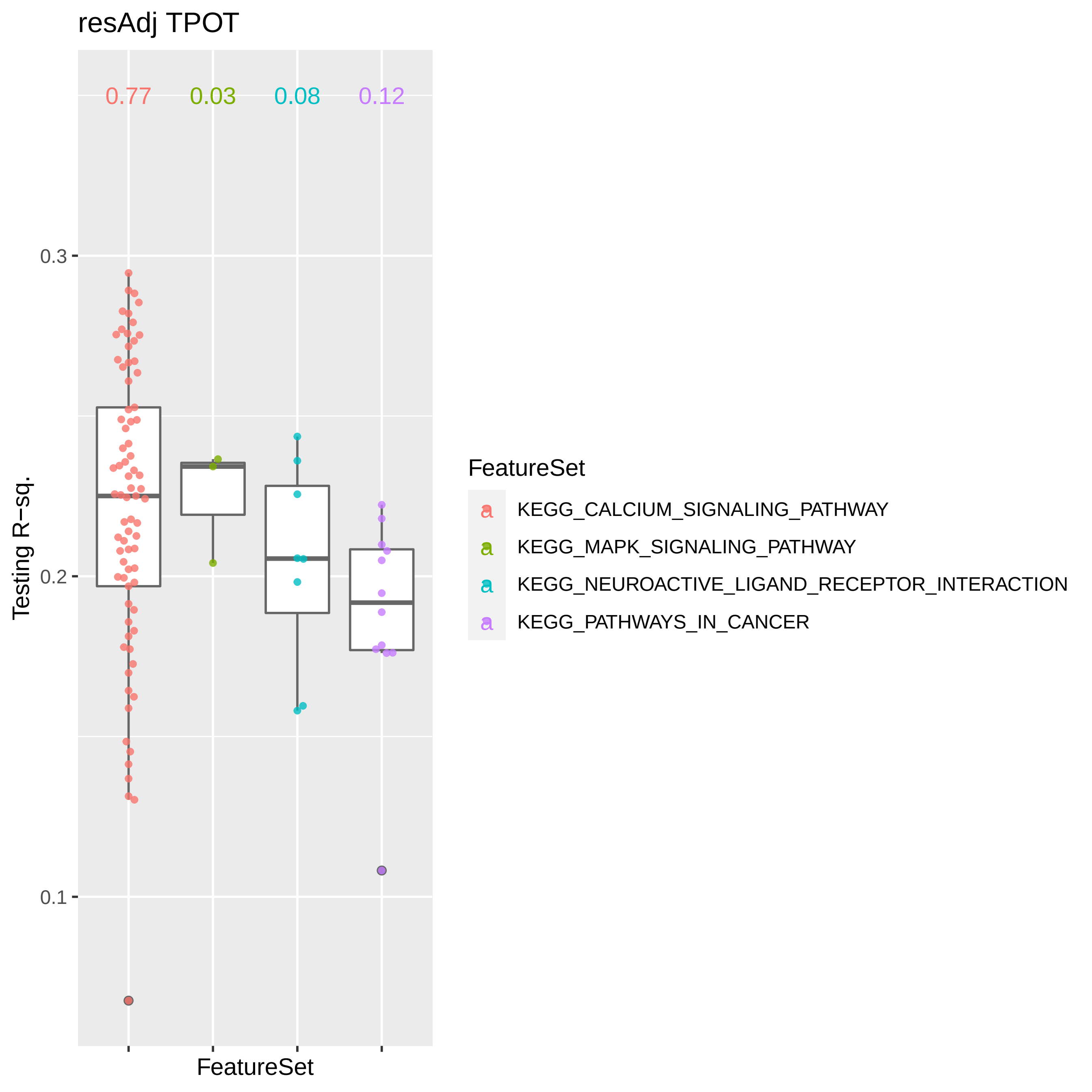

### Additional file 6

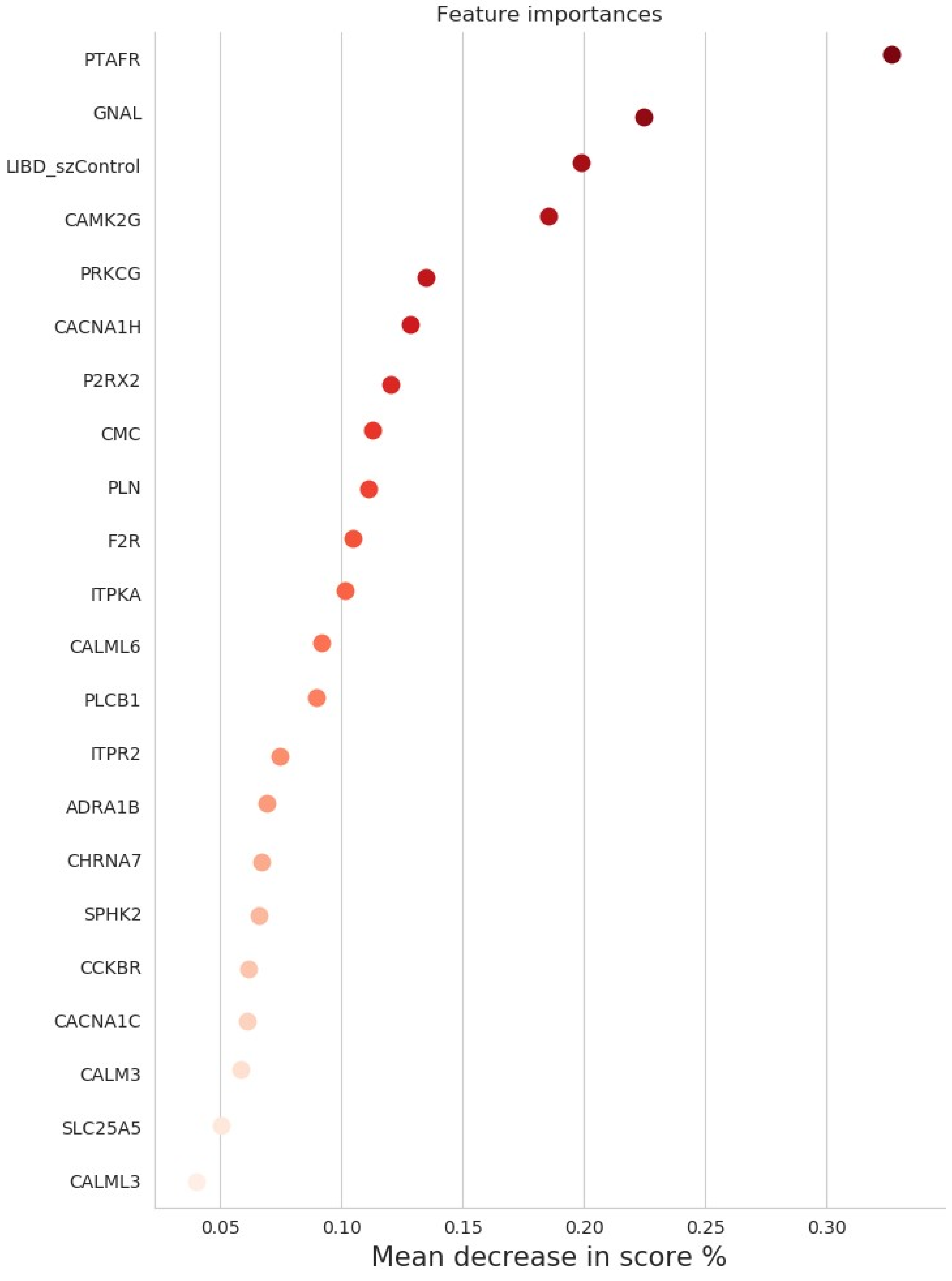

### Additional file 7

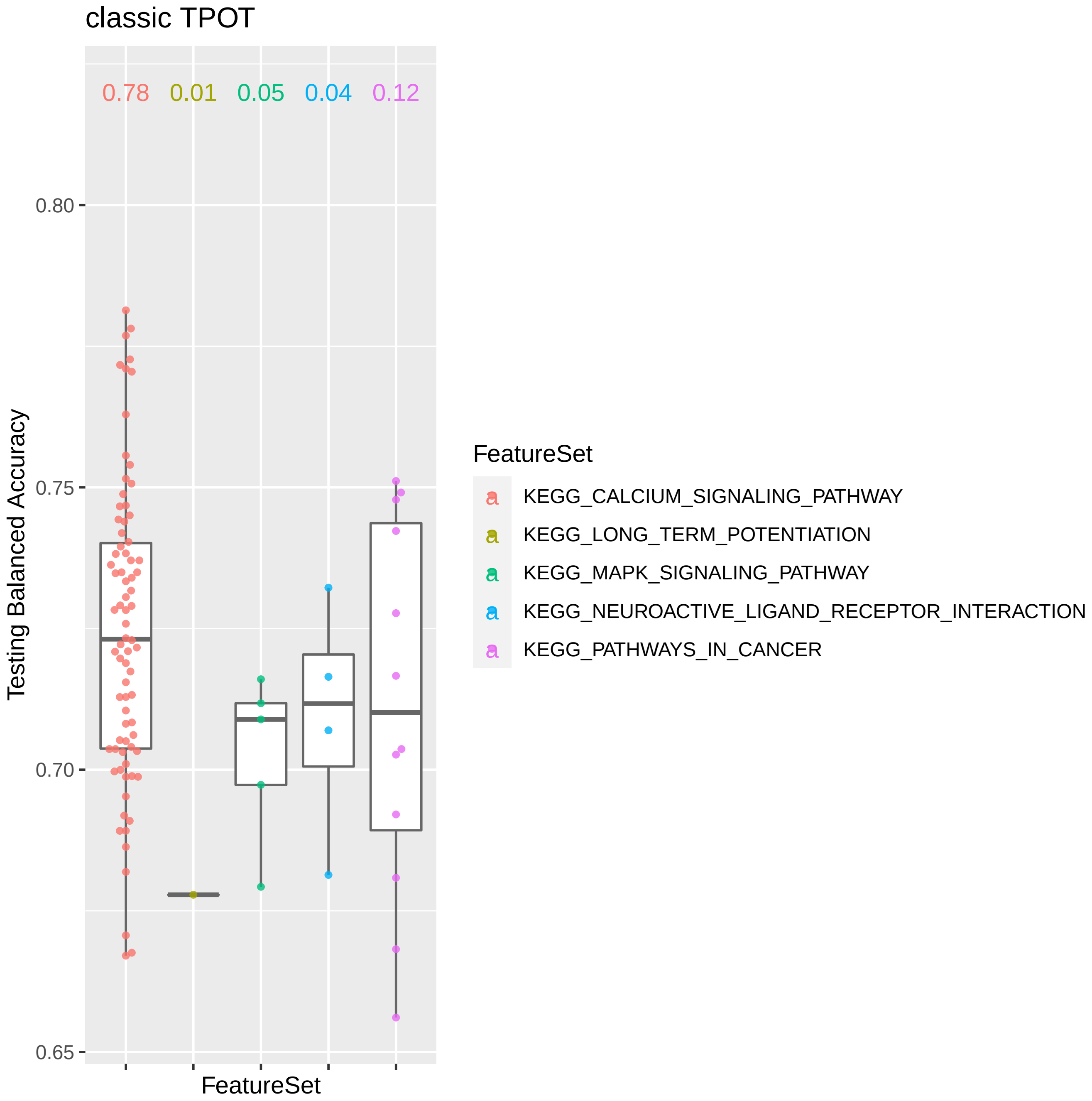

### Additional file 8

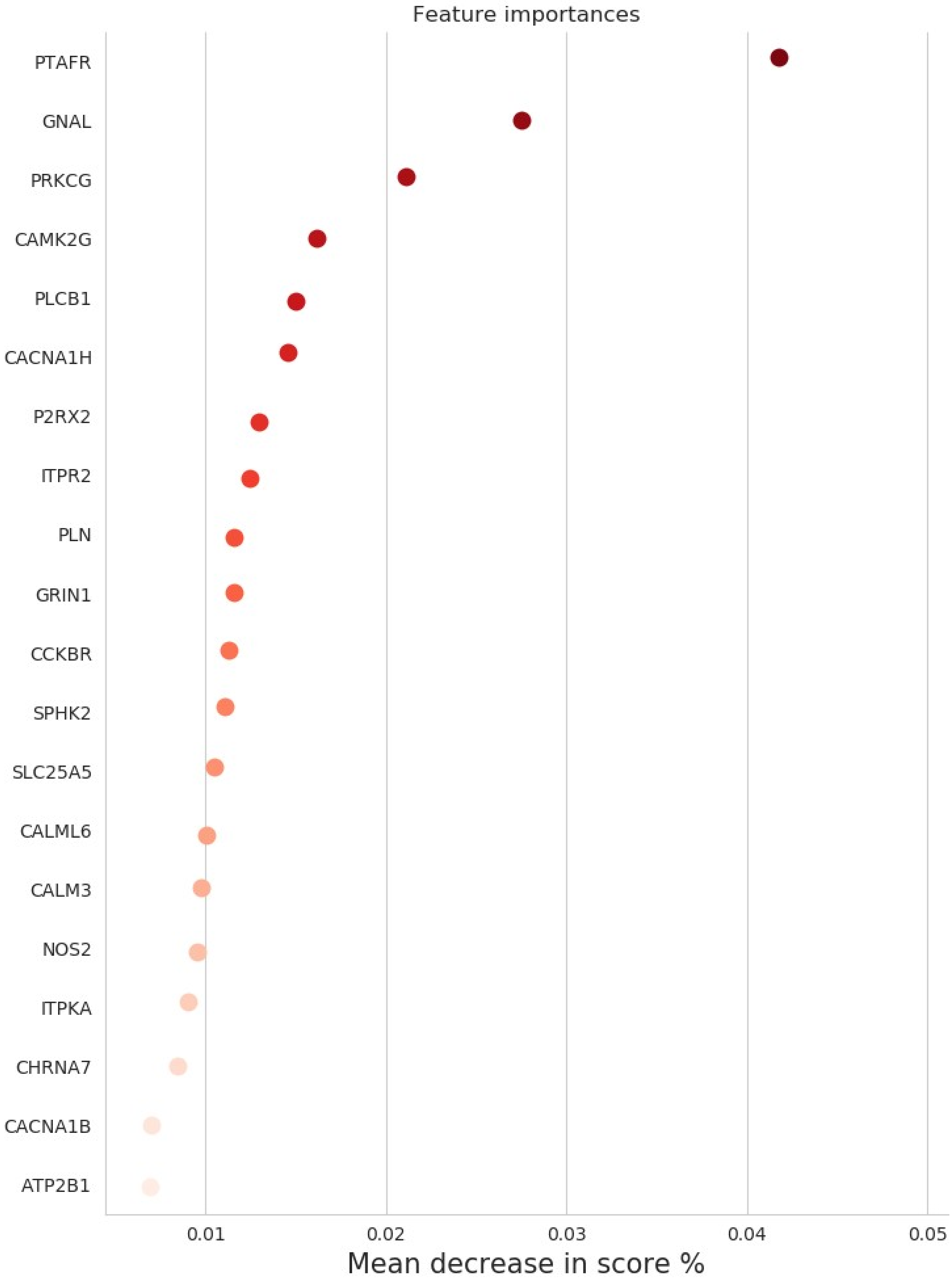
